# Dietary Breadth is Positively Correlated with Venom Complexity in Cone Snails

**DOI:** 10.1101/028860

**Authors:** Mark A Phuong, Gusti N Mahardika, Michael E Alfaro

**Affiliations:** Department of Ecology and Evolutionary Biology, University of California, Los Angeles, CA 90095, USA; Animal Biomedical and Molecular Biology Laboratory, Faculty of Veterinary Medicine, Udayana University Bali, Jl Sesetan-Markisa 6, Denpasar, Bali 80225, Indonesia

**Keywords:** Phylogenetics, comparative biology, venom duct transcriptome

## Abstract

Although diet is believed to be a major factor underlying the evolution of venom, few comparative studies examine both venom composition and diet across a radiation of venomous species. Cone snails within the family, Conidae, comprise more than 700 species of carnivorous marine snails that capture their prey by using a cocktail of venomous neurotoxins (conotoxins or conopeptides). Venom composition across species has been previously hypothesized to be shaped by (a) prey taxonomic class (i.e., worms, molluscs, or fish) and (b) dietary breadth. We tested these hypotheses under a comparative phylogenetic framework using ecological data in conjunction with venom duct transcriptomes sequenced from 12 phylogenetically disparate cone snail species, including 10 vermivores (worm-eating), one molluscivore, and one generalist. We discovered 2223 unique conotoxin precursor peptides that encoded 1864 unique mature toxins across all species, >90% of which are new to this study. In addition, we identified two novel gene superfamilies and 16 novel cysteine frameworks. Each species exhibited unique venom profiles, with venom composition and expression patterns among species dominated by a restricted set of gene superfamilies and mature toxins. In contrast with the dominant paradigm for interpreting Conidae venom evolution, prey taxonomic class did not predict venom composition patterns among species. Our results suggests that cone snails have either evolved species-specific expression patterns likely as a consequence of the rapid evolution of conotoxin genes, or that traditional means of categorizing prey type (i.e., worms, mollusc, or fish) and conotoxins (i.e., by gene superfamily) do not accurately encapsulate evolutionary dynamics between diet and venom composition. We also found a significant positive relationship between dietary breadth and measures of conotoxin complexity. These results indicate that species with more generalized diets tend to have more complex venoms and utilize a greater number of venom genes for prey capture. Whether this increased gene diversity confers an increased capacity for evolutionary change remains to be tested. Overall, our results corroborate the key role of diet in influencing patterns of venom evolution in cone snails and other venomous radiations.

## Introduction

The use of venom for predation has evolved several times across the animal kingdom in organisms such as snakes, snails, and spiders [1]. The majority of venoms consist of complex mixtures of toxic proteins [1–3] and extensive variation in venom composition is documented at nearly all biological scales of study ranging from individuals to species [4–7]. Understanding the forces that shape venom evolution and variation in venom composition among predatory venomous taxa is not only of intrinsic interest to ecological and evolutionary studies [8], but has far reaching implications across several biological disciplines, including drug development in pharmacology and understanding protein structure-function relationships in molecular biology [9,10]. Diet is thought to be a major driver of venom composition patterns because venom is intricately linked to a species’ ability to capture and apprehend prey [8,9]. There are currently two major hypotheses that attempt to explain the impact of diet on broad-scale patterns of venom composition across taxa: (1) prey preference should determine venom components and (2), dietary breadth should be positively correlated with venom complexity [4,11,12]. While both hypotheses are often used interchangeably as evidence for the role of diet in venom evolution (e.g., [11,13,14]), they have separate and distinct predictions on patterns of venom composition among taxa: whereas the former hypothesis predicts the types of venom proteins expected for a given species, the latter hypothesis predicts how many peptides are employed for prey capture.

The idea that prey preference should determine the types of venom proteins employed by a given species is grounded in the logic that natural selection shapes the venom repertoires of species to become more effective at targeting the physiologies of their prey [4,9]. Several studies support this relationship, including correlations between variation in diet and venom components among populations within species [4,15] and functional studies which show that the toxic effects of venoms from different species were maximally effective on their preferred prey [6,16–18]. For example, snake venoms from species that preferentially feed on arthropods were more toxic upon injection into scorpions relative to venom extracted from a species that feeds almost exclusively on vertebrates [19]. However, there are cases where variation in venom composition cannot be attributed to dietary preferences, challenging the generality of this pattern [20,21]. Indeed, gene duplication, positive selection, and protein neofunctionalization are defining features of venom gene evolution [22–25] and these forces work in concert to promote divergence in venom composition among taxa. Given the high evolutionary lability of venom toxins, it is unclear that a relationship between dietary preference and venom composition should be expected.

The second hypothesis on dietary breadth and venom complexity seeks to explain why some species employ more venom proteins than others for prey capture [26]. Under this hypothesis, dietary breadth should be positively correlated with venom complexity because a greater number of venom proteins is necessary to target a wide variety of prey species [11,12]. Although rarely invoked in venom studies, this relationship is explicitly predicted by the niche variation hypothesis, which posits that individuals or populations with wider niches should display greater phenotypic variance [27]. To date, nearly all evidence supporting the impact of dietary breadth in shaping patterns of venom complexity are observational. For example, sea snakes, which mostly feed on fish, have less diverse venoms compared to land snakes, which typically feed on arthropods, reptiles, amphibians, birds, and mammals [11]. In addition, prey specialists tend to have less complex venom compositions compared to generalists [11,28,29]. Despite the apparent signal, these observations have yet to be tested in a phylogenetically controlled and rigorous manner.

Although diet is widely accepted as the dominant force governing venom evolution across disparate venomous taxa [8], few multi-species comparative studies exist that explicitly examine the impact of diet on venom composition patterns across venomous radiations. The majority of studies implicating the prominent role of diet in venom evolution are based on variation in venom composition among populations within species or among closely related species [4,14,19,20,30]. In some cases, broad generalizations on the evolutionary trends of venom and diet are made from the analyses of a few individuals from a single species (e.g, [12,31]). In addition, knowledge on venom composition is often incomplete – most studies are restricted to commonly known gene families [32,33], challenging the generality of previous results given that a substantial proportion of venomous cocktails potentially go unexamined. Without employing a broad and robust comparative phylogenetic approach in conjunction with comprehensive venom data, it is not possible to determine whether previously reported patterns represent general evolutionary trends in venomous taxa or are idiosyncratic phenomena restricted to the particularities of a given study.

Here, we examine the influence of both dietary preference and dietary breadth on venom evolution in cone snails (Family: Conidae), a hyper diverse group of over 700 predatory marine snails that typically prey on either worms, molluscs, or fish using a cocktail of venomous neuropeptides (known as conotoxins or conopeptides) [34,35]. Each species’ venom repertoire is estimated to contain 50-200 peptides, though some recent studies suggest that this number could be in the thousands [31,36,37]. Most conotoxins are typically short in length (~10-30 amino acids long) and target ion channels and neuroreceptors of their prey [38]. Conotoxin precursor peptides typically consist of three regions: (1) a conserved signal sequence containing roughly 20 hydrophobic amino acids that directs the peptide into the secretory pathway, (2) a propeptide region that is cleaved during peptide maturation, and (3) a highly variable mature toxin coding region [35]. Mature toxin coding regions are characterized by their cysteine framework, or the arrangement of cysteine residues often present in mature peptides, which can sometimes provide information on peptide structure and function [32]. Conotoxins are classified into more than 30 gene superfamilies (e.g, A superfamily, M superfamily, etc.) usually based on the similarity of the signal sequence which is conserved within superfamilies across species [32]. Other conotoxins, such as conkunitzins and conopressins, are categorized based on their similarity to proteins from other venomous taxa [35]. Conotoxin gene superfamilies generally do not provide predictive information on peptide function because gene superfamilies can possess conotoxins that target a variety of neurological targets [32].

Both prey preference and dietary breadth have been invoked to explain venom evolution in cone snails [28,29,32,39]. Past studies hypothesized that the broad taxonomic categories of major prey types (i.e., worms, molluscs, or fish) should determine which gene superfamilies are critical venom components for prey capture [31,37,40]. For example, the A superfamily was suggested to not be important for vermivorous (worm-eating) species because it was not detected in the venom duct transcriptome of *Conus miles*, a vermivore [31]. Although few broad comparative studies exist that examine variation in venom composition patterns across cone snail taxa, prey taxonomic class remains the dominant framework by which the evolution of cone snail venom is studied [32,38] and forms the basis for categorization of conotoxins on ConoServer, a molecular database of known conotoxins [32]. Alternatively, the rapid evolution of conotoxin gene superfamilies may facilitate species-specific expression patterns that bear no relationship to whether a species feeds on worms, molluscs, or fish [41]. In terms of dietary breadth, much of the support for a relationship between conotoxin complexity and diet diversity comes from species with diets atypical of cone snails (e.g., highly specialized or extremely broad, [28,29]). Thus, like in other venomous systems, a formal test of this hypothesis has not yet been carried out.

To examine the relationship between venom composition and diet in cone snails, we sequenced the mRNA from the venom duct of 12 cone snails species representing two genera (i.e., *Californiconus* and *Conus*) and spanning nine subgenera within *Conus* (i.e., *Stephanoconus*, *Strategoconus*, *Virroconus*, *Harmoniconus*, *Conus*, *Lividoconus*, *Virgiconus*, *Rhizoconus*, and *Puncticulis*), recovering a broad phylogenetic sampling of species within Conidae [34]. These 12 species consist of 10 vermivores (worm-eaters), one molluscivore, and one generalist that feeds on worms, molluscs, and fish [42]. We describe baseline patterns of venom gene expression across these 12 species and characterize conotoxin composition patterns in the context of a fossil calibrated phylogeny generated from non-venomous, orthologous transcripts. Here, we test two hypotheses: (1) that traditional prey diet categories predict venom composition patterns and (2) that conotoxin complexity is positively correlated with dietary breadth. We analyze ecological data with venom composition patterns under a comparative phylogenetic framework to test the influence of these two hypotheses in shaping patterns of venom evolution.

## Results

We extracted RNA from the venom duct of 12 species (1 individual per species): *Californiconus californicus, Conus arenatus, Conus coronatus, Conus ebraeus, Conus imperialis, Conus lividus, Conus marmoreus, Conus quercinus, Conus rattus, Conus sponsalis, Conus varius,* and *Conus virgo*. Here, we note that *C. sponsalis* refers one lineage of the *C. sponsalis* species complex, where a number of described species are comprised of several, paraphyletic lineages [43]. We use the name *C. sponsalis* to refer to this species complex pending taxonomic revision of this group. We synthesized RNAseq libraries and multiplexed all individuals on a single Illumina HiSeq 2000 lane. We recovered an average of 25.8 million reads per species (Table S1) and assembled transcripts using Trinity [44]. The number of contigs assembled ranged from 28,878 to 88,052, n50 was 609.25 on average, and the total bases assembled ranged from 15MB to 50MB (Table S1).

We used a combination of custom Python scripts, BLAST+, ConoSorter (an algorithm used to identify transcripts that code proteins which share similar properties to known conotoxins), and ConoPrec (a tool used to analyze conopeptide precursors) to identify, filter, and classify conopeptides [45–47]. Through the investigation of conotoxin gene superfamily classifications, we noted several cases where changes in the current naming and classification of gene superfamilies were warranted. Based on signal sequence similarity or protein domain similarity, we reclassified the Divergent_MTFLLLLVSV superfamily as conkunitzins and reclassified the Divergent_MSTLGMTLL superfamily as the N superfamily. We observed that the Divergent_M---L-LTVA superfamily contained several conopeptide precursors with unique and divergent signal sequences. We dissolved this gene superfamily and reclassified it along with all other conopeptides that we were not able to assign into known gene superfamilies.

We used a percent signal sequence identify cut off of 70% to cluster unassigned conopeptides. We assign new names to (1) novel groupings of conotoxin gene superfamilies and (2) groups of conopeptides with similarity to previously characterized conotoxins, but were not given a formal classification. In total, we identified 2223 unique conopeptide precursor sequences encoding 1864 unique mature proteins, 1685 of which are new to this study (Table 1, Table S2). A substantial proportion of these conopeptides were never assembled by Trinity (7.2% - 31% per species, Table S3), but discovered through read mapping and manual reconstruction. These conopeptides span 58 gene superfamilies, nine of which represent gene superfamilies given new names due to reclassification and two of which are newly described (Table S2, Table S4, Fig. S1). Several of these gene superfamilies were recently characterized (e.g. G-like superfamily, SF-mi1 superfamily, etc.), but not given conventional gene superfamily names (i.e. a letter sometimes followed by an Arabic numeral). Although we expanded the membership of these gene superfamilies, we refrained from changing their names pending functional experiments to determine their role in prey apprehension or defense.

**Table 1.** Conotoxin composition and diet for each species analyzed in this study.

| Genus/subgenus | Species | No. of unique conotoxin precursors | No. of unique mature toxins | No. of gene superfamilies | No. of cysteine frameworks | Most abundant gene superfamily and frequency* | Main diet summarized from [29, 42] |
| --- | --- | --- | --- | --- | --- | --- | --- |
| *Puncticulis* | *arenatus* | 326 | 256 | 36 | 31 | O1, (20.3%) | eunicids, nereids, capitellids |
| *Californiconus* | *californicus* | 185 | 164 | 30 | 21 | O1, (20.1%) | molluscs, polychaetes, fish |
| *Virroconus* | *coronatus* | 331 | 286 | 32 | 30 | O1, (19.6%) | eunicids, capitellids |
| *Virroconus* | *ebraeus* | 75 | 69 | 27 | 23 | M, (31.9%) | eunicids, nereids |
| *Stephanoconus* | *imperialis* | 70 | 66 | 20 | 19 | P, (16.7%) | amphinomids |
| *Lividoconus* | *lividus* | 244 | 204 | 31 | 25 | O1, (10.7%) | enteropneusts, terebellids |
| *Conus* | *marmoreus* | 81 | 69 | 14 | 16 | M, (26.1%) | gastropods |
| *Lividoconus* | *quercinus* | 97 | 78 | 25 | 23 | O1, (15.4%) | enteropneusts, sabellids |
| *Rhizoconus* | *rattus* | 102 | 89 | 28 | 30 | con-ikot-ikot, (18%) | eunicids |
| *Harmoniconus* | *sponsalis* | 401 | 338 | 35 | 29 | O1, (28.1%) | eunicids, nereids |
| *Strategoconus* | *varius* | 198 | 168 | 29 | 24 | M, (10.7%) | polychaetes |
| *Virgiconus* | *virgo* | 113 | 78 | 25 | 21 | O1, (24.4%) | terebellids |

\*calculated from no. of unique mature toxins

We identified 70 unique cysteine frameworks across all the conotoxins identified from this study, 16 of which display a novel cysteine motif not yet described from cone snails (Table S5). We report 34 new associations between previously identified cysteine frameworks and gene superfamilies (Table S5). Of particular note, we identified cysteine-free conotoxins from the A and O2 superfamilies, which is in contrast to the cysteine-containing toxins previously described from these groups (Table S5, [35]).

While comparing conotoxins described in this study with the ConoServer database, we identified several discrepancies concerning the species of origin for particular conopeptides. For example, although we did not detect the Qc23a precursor peptide (originally described from *C. quercinus*) in our *C. quercinus* transcriptome, we found a precursor peptide with 100% identity in our *C. imperialis* transcriptome. In another case, nearly every protein (i.e., 70/79 proteins) identified from a recent study on *C. flavidus* [48] had >95 % sequence identity to proteins identified from our *C. lividus* transcriptome. Many of these species mismatches occur between distantly related taxa, where high identity between full precursor peptides is not expected [49]. We hypothesized that several of these discrepancies are cases of species misidentification because we confirmed species identification in this study morphologically and genetically using mitochondrial DNA sequences from the transcriptome. We note these instances in the supplementary for further inquiry (Table S6).

Across all species examined in this study, the number of unique conotoxin precursors ranged from as low as 70 conopeptides in *C. imperialis* to as high as 401 conopeptides in *C. sponsalis* (Table 1). These precursors encoded between 66 to 338 unique mature toxins (also from *C. imperialis* and *C. sponsalis*, respectively, Table 1). We detected only one instance where *C. coronatus* and *C. virgo* expressed the same mature toxin (Co_O2_13, Co_O2_14, and Vi_O2_7, Fig S2). In all other cases, each species expressed a unique repertoire of mature toxins with no overlap between species.

Species varied widely in which gene superfamilies were expressed (Table 1). On average, each species expressed 28 gene superfamilies, with *C. marmoreus* expressing the lowest number of superfamilies (14 superfamilies, Table 1) and *C. arenatus* expressing the highest number of superfamilies (36 superfamilies, Table 1). Only four gene superfamilies were expressed by all the species examined in this study (M, N, O1, and O2, Table S2). These four superfamilies were also the only superfamilies in common amongst the vermivores. The distribution of conopeptides across gene superfamilies per species tended to be skewed, such that > 50% of the conotoxins originated from three to six gene superfamilies (Table S7). The O1 superfamily contained the highest number of mature toxins for seven species, the M superfamily was the most abundant for three species, and the P and con-ikot-ikot superfamilies were each the most abundant for one species (Table 1). Interestingly, the O1, M and con-ikot-ikot superfamilies were also amongst the most abundant conotoxins from a recent study from the transcriptomes of *Conus tribblei* and *Conus lenavati* [41]. The average number of cysteine frameworks found in each species was 24, with *C. arenatus* expressing the highest number of cysteine motifs (31 frameworks, Table 1) and *C. marmoreus* expressing the lowest number of cysteine motifs (16 frameworks, Table 1).

We report a single instance where a premature stop codon interrupts the coding region of an O1 conotoxin expressed by *C. sponsalis* (Sp_O1_79, Fig. S3). The stop codon appears within the signal region and the predicted mature conotoxin is identical to another conotoxin expressed by *C. sponsalis* (Sp_O1_87, Fig. S3). The pseudogenized copy, Sp_O1_79, is more highly expressed than the functional copy, Sp_O1_87, by two orders of magnitude (TPM = 1242.59 and TPM = 11.66, respectively, Fig. S3).

We employed an all-by-all blast approach using the *Lottia gigantea* protein database [50] as our reference to identify 821 putatively orthologous loci suitable for phylogenetic analysis. These loci represent a total of 863,132bp and each species had, on average, 88.3% of the total bases possible in the data matrix. We inferred a maximum likelihood phylogeny in RAxML and generated a time tree using the program r8s with two fossil calibrations from previous studies (Fig. 1, [42,51,52]). The phylogeny was highly resolved with all but two nodes having 100% bootstrap support (Fig. 1).

**Figure 1.**
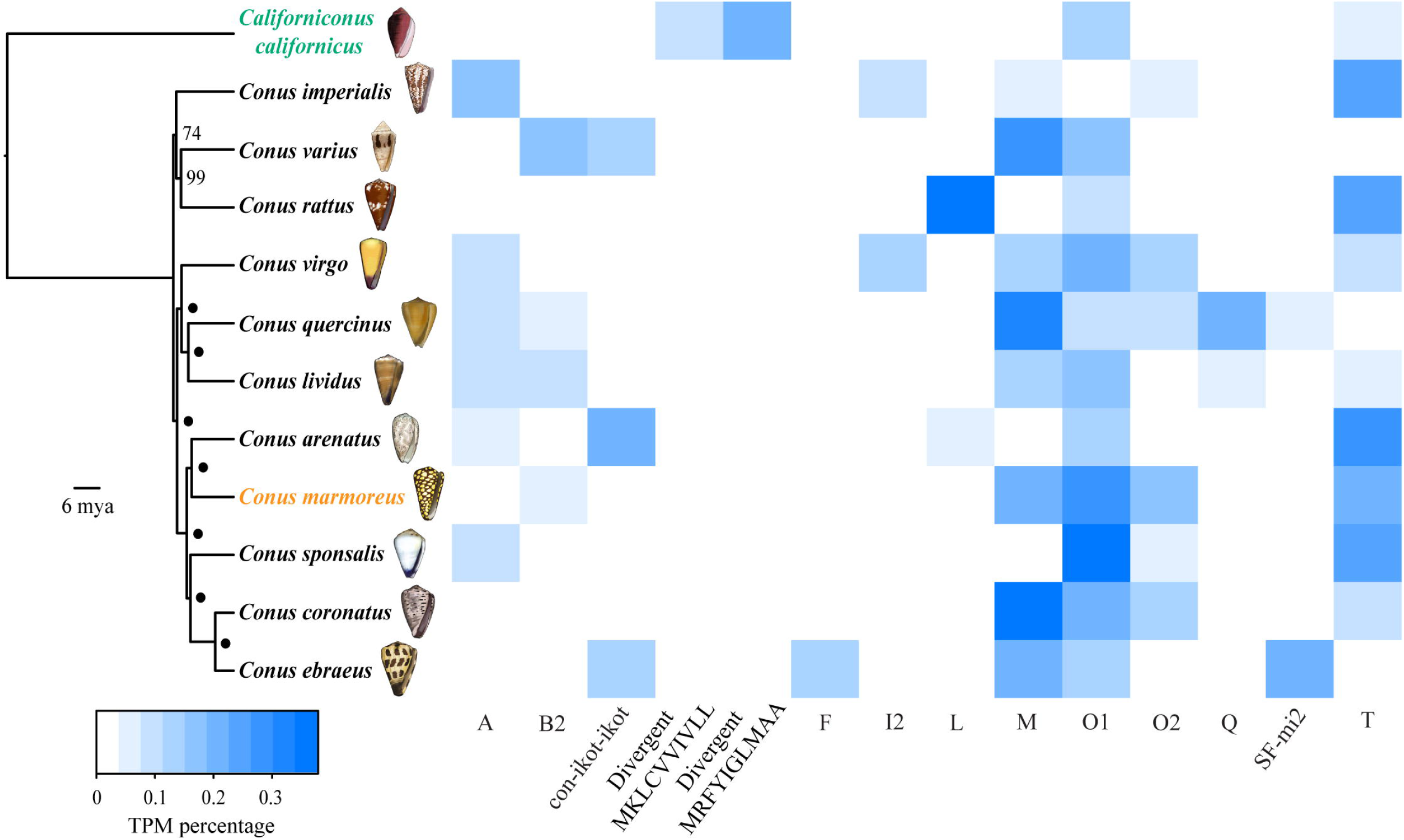
Conotoxin expression in a phylogenetic context. Time-calibrated maximum likelihood phylogeny of Conidae species sequenced in this study generated from 821 loci. Values at nodes represent bootstrap support and • indicates bootstrap support = 100. Tree is rooted with *Californiconus californicus*. Taxa are colored by diet (green = generalist, black = vermivore, orange = molluscivore). Heat map shows relative contribution (measured as percentage of total conotoxin TPM per species) of gene superfamilies that contributed to at least 10% of overall conotoxin expression in at least one species.

We used the RSEM algorithm to generate Transcript Per Million (TPM) values to compare venom duct expression levels between species [53,54]. Total conotoxin expression, or the relative expression of conotoxin genes compared to other genes in the transcriptome, averaged 53% among species and ranged from as low as 26% in *C. californicus* to as high as 70.7% in *C. coronatus* (Table 2). The most highly expressed gene superfamily was the M superfamily for four species, the T superfamily for two species, the O1 superfamily for four species, and the Divergent_MRFYIGLMAA and L superfamilies each being the most abundant for one species (Table 2). On average, the most abundantly expressed gene superfamily represented 28.0% of total conotoxin transcripts (Table 2). In *C. ebraeus*, *C. marmoreus*, *C. sponsalis*, and *C. virgo*, the most abundantly expressed gene superfamily did not contain the most highly expressed mature conotoxin (Table 2). For example, while the most abundant gene superfamily was the M superfamily for *C. ebraeus*, the most highly expressed transcript was Eb_SF-mi2_2, a conotoxin from the SF-mi2 superfamily (Table 2). The average contribution of the highest expressed mature toxin from each species to overall conotoxin expression was 16.1% (Table 2). Conotoxin expression patterns tended to be dominated by a few gene superfamilies and mature conotoxins, such that 2-5 gene superfamilies and 2-23 mature toxins represented more than half of each species’ conotoxin expression levels (Table 2, Table S8). Amongst the most highly expressed gene superfamilies (i.e., representing > 50% of conotoxin expression levels), we did not identify a single superfamily that was shared across all species (Table S8). We identified 14 gene superfamilies with expression levels contributing to at least 10% of overall conotoxin expression in at least one of the species examined in this study, whereas 33 superfamilies never constituted more than 5% of total conotoxin expression in any of the species (Fig. 1, Table S9).

**Table 2.** Conotoxin expression patterns among species.

| Species | Total conotoxin expression | Most highly expressed gene superfamily (frequency) | No. of superfamilies representing > 50% TPM values | Most highly expressed mature toxin (superfamily, frequency) | No. of mature toxins representing > 50% TPM values |
| --- | --- | --- | --- | --- | --- |
| *arenatus* | 57.6% | T (30.2%) | 2 | Ar_T_9 (T, 10.0%) | 11 |
| *californicus* | 26.0% | Divergent_MRFYIGLMAA (20.5%) | 4 | Cl_DivMRFYIGLMAA_6 (Divergent_MRFYIGLMAA, 16.3%) | 10 |
| *coronatus* | 70.7% | M (37.9%) | 2 | Co_M_18 (M, 13.5%) | 11 |
| *ebraeus* | 45.4% | M (21.9%) | 3 | Eb_SF-mi2_2 (SF-mi2, 15.9%) | 5 |
| *imperialis* | 64.3% | T (26.1%) | 3 | Im5.4 (T, 23.2%) | 5 |
| *lividus* | 56.1% | O1 (17.2%) | 5 | Li_O1_25 (O1, 5.2%) | 18 |
| *marmoreus* | 67.9% | O1 (27.3%) | 3 | MaI51 (O2, 17.2%) | 6 |
| *quercinus* | 49.5% | M (32.9%) | 2 | Qc_M_13 (M, 30.5%) | 4 |
| *rattus* | 35.5% | L (36.7%) | 2 | Rt_L_3 (L, 29.3%) | 2 |
| *sponsalis* | 55.7% | O1 (36.7%) | 2 | Sp_A_4 (A, 6.0%) | 23 |
| *varius* | 38.5% | M (26.6%) | 3 | Vr3-SP02 (M, 16.8%) | 5 |
| *virgo* | 68.9% | O1 (21.3%) | 4 | Vi_M_2 (M, 8.9%) | 11 |

We employed the similarity statistic Schoener’s D, a value commonly used to measure niche overlap in diet and/or microhabitat, to quantify the degree of overlap between conotoxin composition among cone snail species with different diets [55]. D values can range from 0 (no overlap) to 1 (complete overlap) [55]. To quantify venom composition similarity, we calculated the D statistic for (a) the percentage of mature toxins belonging to each gene superfamily (referred to as D_mature_) and (b) the percent expression of gene superfamilies (referred to as D_expression_) between all possible pairwise species comparisons. D_mature_ (avg = 0.48, range = 0.28 – 0.7) values on average, were higher than D_expression_ (avg = 0.37, range = 0.09 to 0.68) values (Table S10). To control for phylogenetic signal, we generated residuals from a linear model between both values of D and pairwise phylogenetic distances from the time calibrated phylogeny. We used the residuals in an analysis of variance (ANOVA) to determine whether the distribution of conotoxin overlap values differed depending on whether or not the pairwise species comparisons consisted of (a) a generalist and a vermivore, (b) a molluscivore and a vermivore, or (c) two vermivores. ANOVA results revealed no significant differences between these categories in both D_mature_ (ANOVA, F = 1.69, p > 0.05) and D_expression_ (ANOVA, F = 2.26, p > 0.05, Fig. 2).

**Figure 2.**
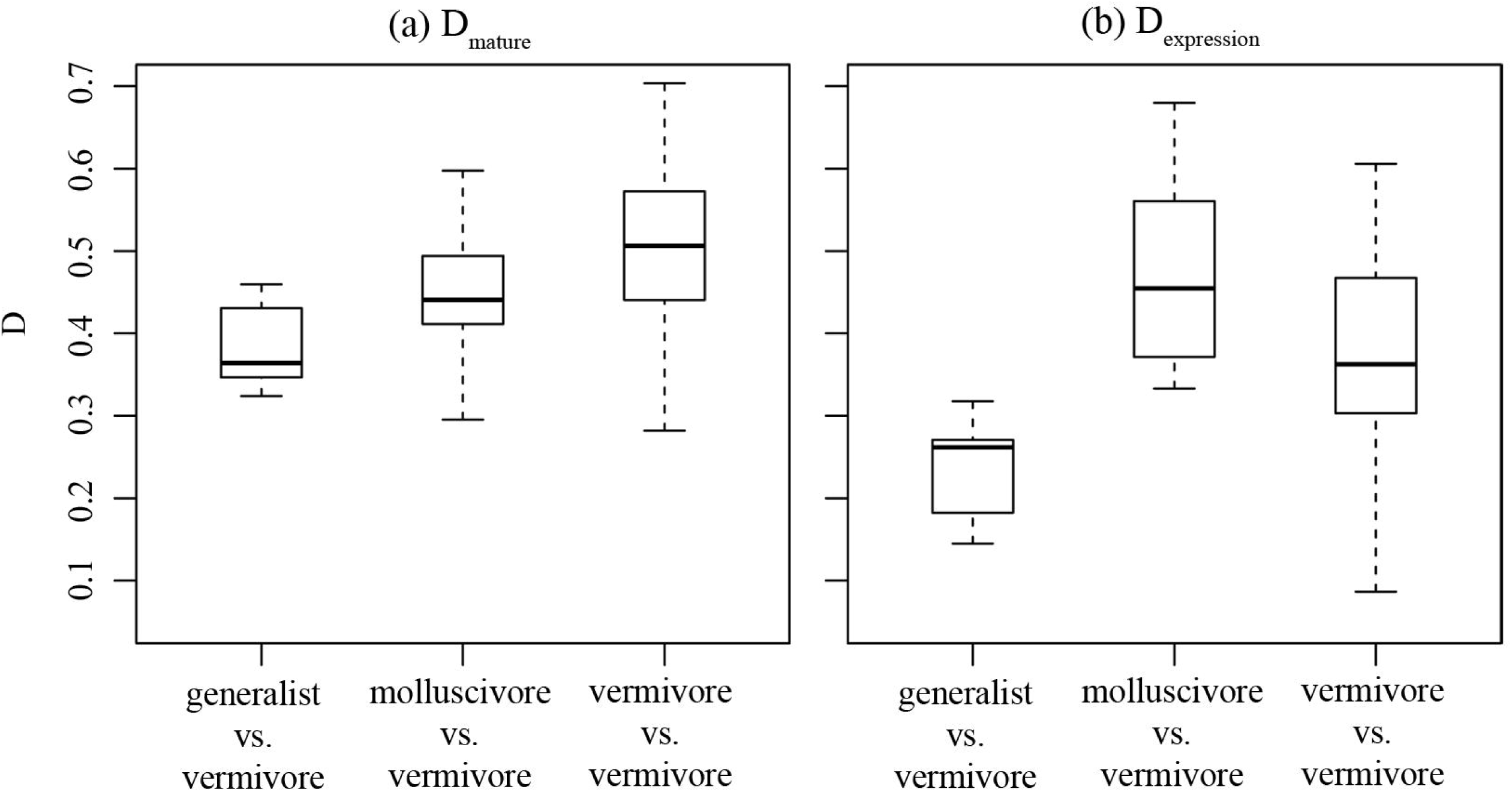
Conotoxin composition similarity between species with different diets. Boxplots showing the distribution of the overlap metric values (D) for conotoxin composition, categorized by whether the species comparison occurred between a generalist and a vermivore, a molluscivore or a vermivore, or two vermivores.

To quantify dietary breadth, we retrieved Shannon diversity index values (H′) representing the diversity of prey species consumed by 10 of the cone snail species in this study that was available in the literature (Table S11, [30,56–61]). To account for phylogenetic non-independence in regression analyses between conotoxin complexity and dietary breadth, we used a phylogenetic generalized least-squares (PGLS) analysis implemented in the caper package within R [62]. We found a significant positive relationship between averaged H′ derived from the literature and the number of mature toxins (PGLS, λ = 1, p < 0.001), the number of gene superfamilies (PGLS, λ = 0.861, p < 0.05), and the number of cysteine frameworks (PGLS, λ = 1, p < 0.05) (Table S12, Fig. 3). The inclusion of *C. californicus* in the PGLS analysis may bias the results because *C. californicus* is often regarded as an atypical member of Conidae due to its extremely broad diet and its distant phylogenetic relationship to the rest of Conidae [29,34]. When removed, relationships remained significant between dietary breadth and the number of mature toxins (PGLS, λ = 0, p < 0.001), the number of gene superfamilies (PGLS, λ = 0, p < 0.05), but not the number of cysteine frameworks (PGLS, λ = 0, p > 0.05) (Table S12).

**Figure 3.**
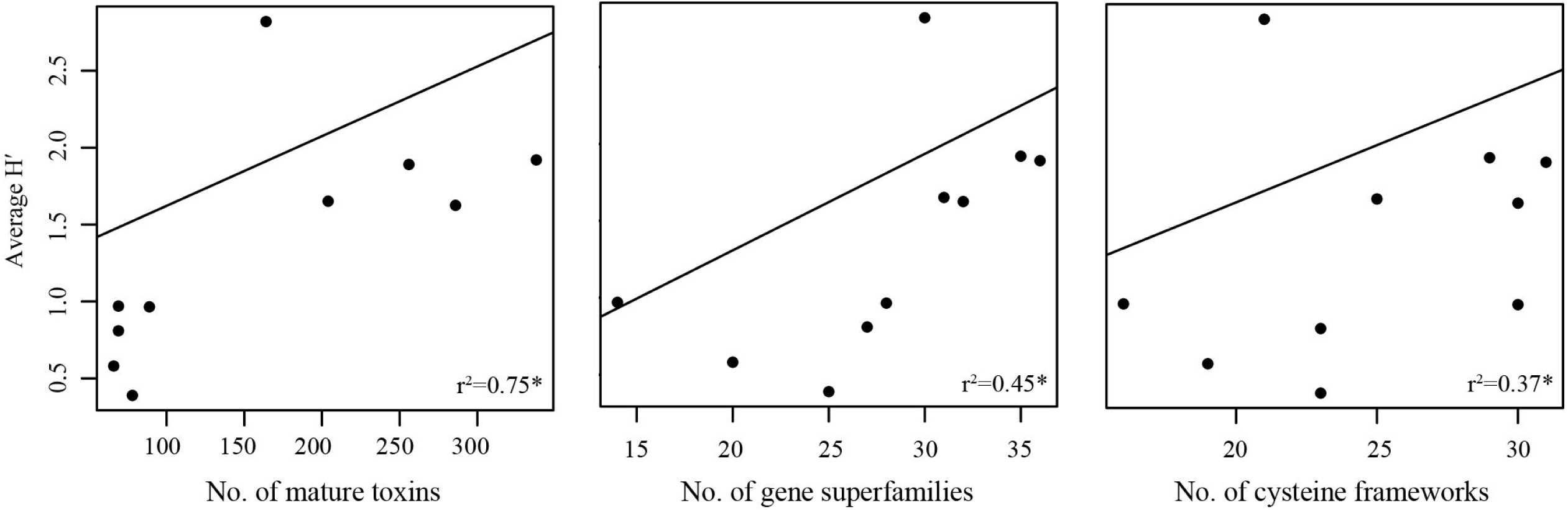
Dietary breadth and conotoxin complexity. Correlations between dietary breadth (Averaged H′) and measures of conotoxin complexity: number of mature toxins, number of gene superfamilies, and number of cysteine frameworks. Graphs are labelled with correlation coefficients. *denotes significant correlation from a PGLS analysis.

## Discussion

### Broad scale patterns of cone snail venoms

Through analysis of venom duct transcriptomes of 12 species spanning a broad phylogenetic distribution within Conidae, we were able to establish baseline characteristics of conotoxin composition and expression patterns in cone snails. For eight species within this study, the total number of predicted mature toxins was within the range of decades-old estimates of 50-200 conotoxins expressed per species (Table 1, [36]). The number of mature toxins for the four remaining species was above 200 (Table 1). Our species-level estimates of conotoxin diversity conflict with a recent study that documented the existence of 3303 conotoxin precursor peptides from the venom duct transcriptome of *Conus episcopatus* [63]. Even when considering the total number of unique conotoxin precursors identified in this study across 12 species (2223 precursors, Table 1, Table S2), this value is still significantly less than what was reported from *C. episcopatus* [63]. Additionally, estimates of conopeptide diversity from our *C. marmoreus* transcriptome (81 unique conotoxin precursors, Table 1) is much lower than estimates from a previous study on the same species (263 unique conotoxin precursors, [37,46]). We hypothesized that these large differences occurred because of the somewhat common practice among cone snail transcriptome studies to identify conopeptides directly from read depth and subsequently to not verify each conotoxin through read mapping [31,37,46,63]. In many cases, unique conopeptide precursors were only supported by a single read [31,37]. These practices produce over-estimates of conopeptide diversity and can lead to erroneous insights on conotoxin variation among species by confounding biological variation with sequencing errors produced by next-generation sequencing platforms [64,65]. Indeed, recent studies have invoked molecular mechanisms such as ‘mRNA messiness’ [31] and ‘RNA editing’ [51] to explain the unexpected abundance of lowly expressed transcript variants likely caused by sequencing errors. Our results echo the sentiments of a previous study emphasizing the importance of carefully and rigorously examining conotoxin sequences generated by new sequencing technologies [66].

Our results confirmed that conotoxin compositions are dominated by a few gene superfamilies and by a few number of transcripts [37,67,68]. This pattern was evident whether we examined conopeptide counts (Table S7) or expression values (Table 2, Table S8). Interestingly, this pattern where venom compositions are dominated by a minority of toxins is also evident in some species of spiders [69] and snakes [11,70,71]. Conticello *et al*. (2001) hypothesized that lowly expressed transcripts, which represent the majority of the conotoxins in cone snail venoms, may not be critical for prey capture and are thus subject to weaker selection pressures that allow for functional divergence over time. These transcripts may provide the substrate for phenotypic novelty as adaptive regimes change, whereby lowly expressed transcripts that have gained new functions become upregulated in response to the environment [67]. It is unclear whether these processes are occurring in cone snail venoms because it is difficult to determine levels of gene expression that are biologically relevant. Further, the strong relationship between dietary breadth and conotoxin diversity documented in this study suggests that the entire complement of toxins is necessary for species to apprehend prey in their environment, indicating that lowly expressed transcripts likely have a significant role in prey capture (Fig 3).

Our estimates of total conotoxin expression in cone snails species from this study were within the range of values estimated from previous work (Table 2, [28,70,72–74]). The level of transcription devoted to conotoxin production in the venom duct may be related to the frequency in which venom is utilized for prey capture, as implicated in [28]. Several laboratory observations report that some cone snail species swallow their prey whole without first injecting venom, though the extent to which this occurs in nature is unknown [56].

Similar to reports from the *de novo* assembly of the *C. episcopatus* venom duct transcriptome [63], Trinity performed poorly in reconstructing all conopeptide precursors reported in this study (Table S3). This may be attributed to the conservation of the signal sequence in precursor peptides, which essentially function as abundant repetitive regions in venom duct RNAseq data. We hypothesize that algorithms devoted to assembly of repetitive regions (e.g., [75,76]) may streamline efforts to characterize Conidae venom duct transcriptomes, which will be the subject of future research efforts.

### Conotoxin composition and diet

Contrary to common wisdom derived from many previous studies of cone snail venom, our results show unequivocally that prey dietary class performs poorly in predicting conotoxin composition patterns among cone snail species. We did not detect a single gene superfamily that separated vermivores from the molluscivore or the generalist (Table S2, Fig. 2); rather, the defining feature across cone snail venoms was that every species, regardless of diet category, expressed a unique repertoire of conopeptides and gene superfamilies at different magnitudes (Table S2, Table 2, Fig. 1). The uniqueness of each species’ conotoxin composition is underscored by moderate to low D values (Table S10) and no apparent differences in the distribution of D values among species that do or do not share the same diet class (Fig. 2). Unfortunately, we were not able to obtain venom duct RNAseq data for several molluscivores and piscivores, but our results are in agreement with a meta-analysis indicating that there is no gene superfamily that there is exclusive to any of the traditionally recognized diet types [77].

The poor predictive power of diet class on venom composition patterns may be due to several factors. Estimated rates of gene duplication and nonsynonymous substitution rates for conotoxin genes are the highest across metazoans [24,78] and these extraordinary rates of molecular evolution may be more likely to promote divergence rather than convergence in venom composition, as documented in this study. Alternatively, the lack of predictive power may be in part, due to how categories are constructed for venom components and diet classes. Prey specialization exists at the level of protein function, but it is known that conotoxins targeting similar neurological targets can evolve convergently in several gene superfamilies [32,35]. Therefore, characterizing conotoxin composition patterns by gene superfamilies does not fully measure functional similarities and differences in conotoxins among species that share similar diets. Prey classes may not explain conotoxin composition patterns potentially due to an over generalization of the diversity present within the vermivorous category. Vermivory, as traditionally used in Conidae studies, is broadly defined and includes a wide variety of taxa that represent hemichordates, echiurans, and several polychaete families that diverged ≥ 400 million years ago [42,79]. Future studies that account for the taxonomic breadth of worms and the functional diversity of conotoxins may better predict patterns of venom composition among species.

We found strong support for a highly significant, positive relationship between venom composition complexity and dietary breadth across cone snails (Fig. 3), corroborating hypotheses that were made from species that represent the extremes of the dietary breadth spectrum in Conidae [28,29]. The total number of mature toxins provided the strongest predictor of this relationship, possibly because mature toxin diversity better encapsulates functional diversity in cone snail venoms. The relatively weaker correlations documented for gene superfamilies and cysteine frameworks support the notion that these traditional means of classifying conotoxins are not simple or direct correlates of conotoxin function (Fig. 3, Table S12 [66,80]). The positive relationship between dietary niche breadth and venom composition complexity corroborates the niche variation hypothesis, suggesting that diverse venoms are required to subdue diverse prey. Although the majority of studies invoking this hypothesis focus on population level trait variance within species (e.g, [81]), our results extend the generality of this hypothesis to species-level patterns of conotoxin complexity across cone snails. These results also align with observations on venom composition complexity in snakes [11,12,33], suggesting that dietary breadth may explain evolutionary trends in venom complexity across radiations of venomous taxa.

At the population level, cone snails follow the predictions supported by the niche variation hypothesis, such that snail populations with a greater dietary breadth show greater population-level allelic diversity in some conotoxin loci [30,82]. How these population-level patterns translate into species-level properties remains unclear. Although conotoxin allelic diversity is higher in populations with greater dietary breadth, each individual within the population possesses only a small subset of the available alleles per locus because these alleles belong to individual loci and are not separate genes [30,82]. Presumably, gene products encoded by these loci are effective at paralyzing diverse prey and toxins encoded by variants at these loci may be more effective if expressed in tandem [82]. Given the exceptional rates of gene duplication estimated within this group [24], gene duplication presents a mechanism by which allelic variants previously restricted to a single conotoxin locus can ultimately evolve to be expressed simultaneously through duplication, potentially leading to species-level patterns of venom composition complexity and dietary breadth.

What are the evolutionary consequences of increased dietary breadth? Our results imply that dietary breadth plays a role in determining how many conotoxins are utilized for prey capture. Consequently, this also determines the number of genes available for forces such as mutation, selection, and drift to generate novel functions for adaptation. Higher gene diversity is thought to provide increased opportunities for novel phenotypes to arise [83], potentially shaping a lineage’s evolvability, or a lineage’s capacity to evolve in response to their environment [84]. Therefore, greater dietary breadth (leading to higher gene diversity) may signify a larger potential for lineages to diversify, potentially influencing patterns of species diversification in cone snails. Although venom is viewed as a key innovation and thought to play a major role in the evolutionary diversification of venomous taxa [1,85], the interplay between diet and venom on patterns of lineage diversification is rarely tested explicitly. High resolution molecular phylogenies of venomous taxa and comprehensive venom composition data that can now be rapidly obtained using new sequencing technologies will provide the necessary datasets to facilitate an examination of the evolutionary dynamics of diet, venom, and speciation over long evolutionary time-scales.

A recent study showed that cone snail species may inject separate suites of conopeptides for predation and defense, and that defensive venoms are produced in the proximal region of the venom duct while predatory venoms are produced in the distal region [86]. Because we generated our venom duct RNAseq data from the entire length of the organ, our results may be confounded between these two distinct ecological roles that venom performs. However, it is unclear how broad this pattern is throughout cone snails. In many cases, functional work has shown that venoms extracted from different regions of the venom duct were able to successfully paralyze prey [16,17]. In addition, the functional roles of conotoxins that are relevant to each species’ ecology are poorly understood. Conotoxin function is typically determined by assays in mice or vertebrate neuronal cells which are not representative of the intended targets of conotoxins [35]; as previously asserted in snake systems [18,19] over-interpretation of these results can lead to misleading or conflicting inferences. For example, a δ-conotoxin (a type of conotoxin thought to be critical for fish-hunting) isolated from the vermivore, *Conus susturatus*, was implicated as a defensive toxin against fish predators due to its effects on vertebrate sodium ion channels and its proximal expression in the venom duct, where defensive toxins are thought to be synthesized [87]. However, behavioral observations from *Conus tessulatus*, a vermivore in the same subgenus that also expresses a similar δ-conotoxin, was shown to prey on fish on occasion through venom injection [88]. These results emphasize the necessity of examining conotoxins in the context of each species’ ecology to accurately understand the natural history of cone snails.

### Conclusion

In contrast to the most widely accepted hypothesis of cone snail venom evolution, diet class did not predict patterns of venom composition among cone snails. These results suggest either (a) the fast rates of venom evolution drive rapid divergence of conotoxin composition that bear no relationship to prey taxonomic class, or (b) current ways of categorizing both prey species (i.e., worm, mollusc, fish) and conotoxins (i.e., gene superfamily) fail to accurately reflect evolutionary interactions between dietary specialization and venom function. Therefore, future studies placing more emphasis on the taxonomic breadth of cone snail prey and conotoxin function on prey capture may better encapsulate the impact of diet on cone snail venom evolution. In addition, our results highlight the importance of dietary breadth in shaping species-level venom complexity patterns among cone snails. To our knowledge, this relationship is rarely tested quantitatively across venomous radiations despite its potential to explain variation in venom complexity as demonstrated here. While our results show that species with broad diets tend to have more diverse venoms, the evolutionary consequences of this tendency remains unclear. What is certain is that selective pressures driven by diet plays a major role in shaping evolutionary patterns in venom across cone snails and other venomous taxa.

## Methods

### Sampling and sequencing

We collected one individual from 11 species of *Conus* from Bali, Indonesia (*C. arenatus*, *C. coronatus*, *C. ebraeus*, *C. imperialis*, *C. lividus*, *C. marmoreus*, *C. quercinus*, *C. rattus*, *C. sponsalis*, *C. varius*, *C. virgo*) and WF Gilly provided 1 *C. californicus* species from Monterey Bay, California. We immediately placed dissected venom ducts from live snails in RNALater and stored samples in a 4°C refrigerator until they could be placed in a -20°C freezer within 2 weeks of collection. We isolated RNA using TRIzol reagent (Invitrogen, USA) and purified the sample using a Qiagen RNeasy Mini Kit. We extracted RNA from the entire venom duct, or along sections of the venom duct if it was particularly long because venom composition is known to change along the length of the duct in some species [89]. We used Bioanalyzer traces to assess total RNA quality and to determine suitability for sequencing. We constructed cDNA libraries by using the TruSeq RNA Sample Prep Kit to recover mRNA via Poly-A selection, synthesize cDNA, ligate adapters, and barcode samples. We sequenced all 12 samples on a single Illumina HiSeq 2000 lane with 100-bp paired-end reads.

### Assembly

During initial attempts to assemble transcripts in Trinity, we were not able to assemble known transcripts present in the sequencing data, potentially due to the repetitiveness and high sequence complexity of venom transcripts. To circumvent this issue, we employed an iterative assembly approach. For each iteration, we trimmed adapters and low quality bases using Trimmomatic [90], merged reads using FLASh [91], and assembled transcripts using Trinity [44]. During the first assembly iteration, we assembled a 0.1% random subset of the total reads for each sample. Then, we used blastx to identify transcripts with similarity (evalue = 1-e10) to known conotoxin genes listed on ConoServer. We used bowtie2 [92] to align and identify reads that matched to those putative venom transcripts. For the second iteration, we assembled reads from the 0.1% subset that did not align to the venom transcripts from the first iteration. Then, we identified additional putative venom transcripts from the contigs generated. For the final iteration, we assembled reads from the full dataset that did not align to venom transcripts identified from the first two iterations and identified additional contigs that shared similarity to conotoxins.

### Conotoxin identification

We used Conosorter to identify novel venom transcripts and took a conservative approach towards accepting venom transcripts because ConoSorter has a tendency to over-classify sequences. For example, the recently discovered Y2 superfamily identified through ConoSorter is actually molluscan insulin [93]. We used ConoSorter to analyze transcripts with TPM values > 1000 and retained conotoxins that (1) had all three conotoxin regions (i.e., signal region, propetide region, and mature toxin coding region) and (2) a precursor protein length > 38 and < 200 (boundaries were generated based from the empirical length distribution of conotoxin proteins identified from this study). We used blastx to query the novel venom transcripts for similar sequences in every transcriptome in our dataset and also against the ConoServer database. We retained sequences if similar signal regions could be found in the transcriptomes of other species or in ConoServer. We removed transcripts if they produced erroneous blast results (e.g., best-scoring transcript in a different species’ transcriptome produced a protein with several stop codons), suggesting that the novel conotoxin identified by ConoSorter may have been in an incorrect reading frame.

To generate a venom gene reference for each species, we combined all venom transcripts from each assembly iteration and transcripts identified through ConoSorter. We used Python scripts to remove transcripts with redundant proteins and transcripts belonging to the incorrect species. We identified several cases where highly expressed transcripts in one species could be found at low expression levels in some, or in all of the other species sequenced. We note that cross-contamination across every single sample is unlikely, given that these samples were prepared in different sets and were pooled just before sequencing. We hypothesized that this phenomenon occurred due to cluster misidentification during sequencing, potentially due to high sequence similarity of conotoxin transcript signal sequences. We used blastp to identify and remove transcripts from species that had high identity (>95%) in the protein coding region to another species. Here, we do not expect >95% identity across the entire conotoxin precursor protein across the taxa in this study, given the exceptionally high nonsynonymous substitution rates estimated in venom genes from Conidae [37]. We chose the transcript from the species that had the highest coverage (estimated using bowtie2) to be the true transcript.

To reassemble transcripts that were incomplete (missing start or stop codon), we used an approach called Assembly by Reduced Complexity (ARC, https://github.com/ibest/ARC). ARC is a pipeline that allows for *de novo* assembly of specific targets by only assembling reads that map to reference targets. We removed venom transcripts that could not be reassembled, or were not full length (included a start and stop codon) after a maximum of three ARC iterations. Then, we used bowtie2 to map reads to all venom genes to verify the nucleotide sequence of each putative conotoxin. We removed sequences that did not have reads aligning to the entire length of the transcript.

Through mapping, we identified sequence polymorphisms in the conotoxin transcripts. In some cases, these polymorphisms represented allelic differences and we generated a separate conotoxin sequence if the sequence translated into a unique precursor peptide. In other cases, the polymorphisms represented completely distinct conotoxin transcripts that were never assembled, but partially mapped to the existing reference. We assembled these conotoxin transcripts by manually aligning representative reads in Geneious and verified each sequence through additional read mapping. To check for chimeric sequences, we generated 80bp fragments every 20bp along the length of each transcript and searched for the existence of these fragments directly from read depth for sequences that had > 30X coverage. We manually examined sequences flagged by this filter and removed sequences if necessary. We used ConoPrec to remove sequences that did not have a clearly defined signal sequence. Finally, we manually inspected all venom genes to identify any unusual conotoxin transcripts.

### Conotoxin classification

We employed several approaches to classify conotoxins into gene superfamilies. First, we compared conotoxin transcripts to sequences from the ConoServer database using a blastx search and assigned transcripts to gene superfamilies using the best-scoring hit. For transcripts that did not have a blast hit, we compared signal sequences using blastp against the ConoServer database and classified transcripts to gene superfamilies that had a percent signal sequence similarity > 76%, a threshold used in a previous study [39]. We noted that the Divergent_M---L-LTVA superfamily was composed of several transcripts with unique signal sequences. With members of this superfamily along with all other unclassified transcripts, we aligned signal sequences and generated a pairwise distance matrix in Geneious (Biomatters, Auckland, New Zealand). Then, we used a custom Python script to cluster conopeptides that shared a percent signal sequence similarity > 70%. We derived this threshold empirically to minimize the number of clusters, yet still represent salient differences among clusters (e.g., similar cysteine frameworks). We provided names for novel, reclassified, and unclassified superfamilies with five letters representing the first five amino acids that the majority of their constituent sequences shared.

To provide names for conotoxin precursors identified in this study, we followed the naming conventions similar to [41]. Briefly, we named each conotoxin with the following: two letters to denote the species, the gene superfamily name, and a number denoting the order of discovery within the gene superfamily for that species. These fields are separated with an underscore. We did not provide new names for previously identified conotoxins unless there was evidence of species misidentification in previous work.

### Phylogeny inference

For all transcripts not classified as conotoxins, we used CAP3 [94] to reduce redundancy and annotated the transcriptomes using blastx against the *Lottia gignatea* (owl limpet) protein database [50]. To identify putatively orthologous loci for phylogenetic reconstruction, we employed a reciprocal blast approach via blastx and tblastx between each species’ transcriptome and the *L. gigantea* database. We retained loci that had at least 10 species represented. For each of these loci, we also considered other contigs within each species’ transcriptome that was annotated with the same protein, but spanned a non-overlapping portion because transcriptomes are often fragmentary. We created alignments using MAFFT [95] and manually inspected each locus in Geneious. We used a custom Python script to calculate uncorrected patristic distances between all possible pairwise comparisons of the taxa in this study. For each comparison, we removed loci with patristic distances greater than two standard deviations away from the mean to remove potential paralogous sequences. We concatenated all loci and inferred a phylogeny using RAxML under a GTRGAMMA model of sequence evolution with 100 bootstrap replicates [35], rooting our tree using *C. californicus* based on previous phylogenetic hypotheses [21].

We dated the maximum likelihood phylogeny generated from RAxML using the program r8s with two fossil calibrations: a fixed rate of 55 my (million years) representing the origin of cone snails in the fossil record at the root of the tree [51], and a minimum constrained age of 11my (the earliest date showing fossil evidence of both *C. lividus* and *C. quercinus*) at the node representing the ancestor of *C. lividius* and *C. quercinus* [42]. The inclusion of *C. californicus* in this study allowed us to place the fossil calibration at the root because this node possibly represents the ancestor to all Conidae [34].

### Conotoxin expression

We removed transcripts with significant homology to the mtDNA genome of *Conus consors* and the *L. gigantea* non-coding RNA database via blastn to remove potential biases associated with quantifying venom expression. To normalize read counts, we used the RSEM algorithm to map reads with bowtie2 and generate TPM values.

### Diet and conotoxin composition overlap

To calculate overlap in conotoxin composition among species, we employed Schoener’s D statistic [55]: 1

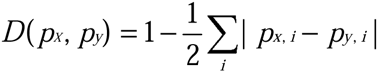

where *p_x_* and *p_y_* represent the frequencies for species x and species y for the ith category. D ranges from 0 (no niche overlap) to 1 (niches are identical). We calculated D by using (a) the percentage of mature conotoxins belonging to each gene superfamily (D_mature_) and (b) the percentage of overall conotoxin expression levels for each gene superfamily (D_expression_) for all possible pairwise species comparisons. We categorized each comparison as either occurring between (a) a generalist and a vermivore, (b) a molluscivore and a vermivore, or (c) between two vermivores. To control for phylogenetic signal, we generated a pairwise phylogenetic distance matrix from the time-calibrated phylogeny using the R package picante. Then, we calculated residuals from a linear regression model between phylogenetic distance and both D values. We used the residuals in an ANOVA to determine whether the distribution of overlap values between the species comparisons were significantly different based on diet.

### Dietary breadth and conotoxin complexity

We obtained dietary breadth measurements estimated by Shannon’s diversity index values (H′) for 10 species from primary literature [30,56–61]. We retrieved H′ values directly from these past studies or calculated them if necessary. When calculating H′, we ignored categories that appeared to represent an amalgamation of several unidentified taxa. We only included H′ values that were calculated from at least 5 individuals where the prey item could be identified to the genus level. We generated average H′ for each species rather than recalculate H′ values for the total number of prey taxa that each species can consume, because each cone snail species preyed on a different set of taxa depending on the geographic locality and what congeners were present [56–60].

We quantified conotoxin composition diversity as either the (a) number of mature toxins, (b) number of gene superfamilies, or (c) number of cysteine frameworks. We correlated these values with averaged values of H′ in a PGLS analysis implemented in the R package caper. When performing regression analyses, the PGLS function in caper incorporates a covariance matrix by using branch lengths from an ultrametric phylogeny and assuming a Brownian motion model of trait evolution [62]. We executed these analyses with and without *C. californicus*.

## Acknowledgements

We thank DST Hariyanto, MBAP Putra, MKAA Putra, and the staff at the Indonesian Biodiversity Research Center in Denpasar, Bali for assistance in the field; the community at the Evolutionary Genetics Lab at UC Berkeley for laboratory support; and J Chang, M Lim, E McCartney-Melstad, WF Gilly, the J McGuire lab group, and the UCLA next-generation sequencing working group for insightful comments on earlier versions of this manuscript. This work was supported by a research grant from the London Malacological Society, a Grants-in-Aid of research from Sigma Xi, a research grant from the Ecology and Evolutionary Biology department at UCLA, the Lerner Gray Fund for Marine Research from the American Museum of Natural History, an Edwin W. Pauley Fellowship from UCLA, a Fulbright Fellowship, and a NSF Graduate Research Fellowship awarded to MAP. This work used the Vincent J. Coates Genomics Sequencing Laboratory at UC Berkeley, supported by NIH S10 Instrumentation Grants S10RR029668 and S10RR027303. We thank the Indonesian Ministry of State for Research and Technology (RISTEK, permit number 277/SIP/FRP/SM/VIII/2013) for providing permission to conduct fieldwork in Bali. The *C. californicus* specimen was collected under a California Department of Fish and Wildlife collecting permit granted to WF Gily (SC-6426). We thank C. Brown for illustrating the images in Figure 1.

## Author’s contributions

MAP designed the study, conducted the field and laboratory work, carried out the bioinformatic analyses, and drafted the manuscript. GNM participated in field work. All authors read, reviewed, and approved the final manuscript.

## Data availability

Raw read data are available at the National Center for Biotechnology Information Sequence Read Archive (Accession Numbers SRX1323883-SRX1323894). All conotoxin transcript sequences will be uploaded to ConoServer. All scripts and final datasets used for analyses will be uploaded onto Dryad.

**Captions:**

**Figure S1. New conotoxin gene superfamilies described in this study.** Signal sequences are underlined, mature toxin regions are bolded, and cysteines within the mature toxin region are highlighted. Ar = *C. arenatus*, Co = *C. coronatus*, Im = *C. imperialis*, Li = *C. lividus*, Qc = *C. quercinus*, Rt = *C. rattus*, Sp = *C. sponsalis*, Vi = *C. virgo*.

**Figure S2. Identical mature toxins from two different species.** Full conopeptide precursor sequences of identical mature toxins expressed in more than one species. Signal sequences are underlined, mature toxin regions are bolded, and cysteines within the mature toxin region are highlighted. Co = *C. coronatus*, Vi = *C. virgo*.

**Figure S3. Pseudogene identified from *C. sponsalis*.** Full precursor peptide sequences from the functional and pseudogenized copy of a mature toxin identified from *C. sponsalis* (Sp = *C. sponsalis*). TPM values are shown, signal sequences are underlined, mature toxin regions are bolded, and cysteines within the mature toxin region are highlighted.

**Table S1. Sequencing and assembly statistics for each species.** Values were calculated using all transcripts assembled from all iterations of Trinity.

**Table S2. Conotoxin diversity by gene family for each species sequenced in this study.** Each value represents the number of unique predicted mature toxins. Values in parentheses represent the number of unique precursor peptides.

**Table S3. Comparison between final conotoxin dataset and conotoxins assembled through three iterations of Trinity.** Number and percent of unique precursor peptides from the final dataset that had at least 95% sequence identity to transcripts assembled by trinity and that matched to (a) 90%, (b) 95%, or (c) 99% of the total precursor sequence length.

**Table S4. Conopeptide gene superfamilies that were reclassified and provided new names in this study.**

**Table S5. Cysteine frameworks identified in this study for each gene superfamily.** * indicates novel cysteine framework discovered in this study. ^a^ indicates a cysteine framework found in cone snail venoms, but not yet described from this gene superfamily.

**Table S6. Conopeptides sequenced in this study with 100% protein sequence identity to a different species on ConoServer.** Matches ≥95% are shown for conopeptides described *Conus flavidus* to highlight similarities with the *C. lividus* transcriptome from this study.

**Table S7. Gene superfamilies that represent > 50% of the conopeptides identified per species.**

**Table S8. Gene superfamilies and conotoxins representing major components of each species’ venom expression levels.** Divergent is abbreviated as Div.

**Table S9. Relative contribution of each gene superfamily to conotoxin expression.** Values calculated as % conotoxin TPM (superfamily TPM/total conotoxin TPM)

**Table S10. Conotoxin composition overlap values (D).** Values calculated by (a) the frequency of conopeptide membership to different gene superfamilies (b) the percent expression level of each gene superfamily.

**Table S11. Dietary breadth, measured by Shannon-Wiener’s index (H′), for 10 cone snail species sequenced in this study.**

**Table S12. Results from Phylogenetic Generalized Least Squares (PGLS) analysis assessing the relationship between prey breadth (H’) and conotoxin complexity.**

## References

1. Fry BG, Roelants K, Champagne DE, Scheib H, Tyndall JDA, King GF, et al. The toxicogenomic multiverse: convergent recruitment of proteins into animal venoms. Annu Rev Genomics Hum Genet. 2009;10: 483–511. doi:10.1146/annurev.genom.9.081307.164356

2. Olivera BM. Conus venom peptides: reflections from the biology of clades and species. Annu Rev Ecol Syst. 2002;33: 25–47. doi:10.1146/annurev.ecolsys.33.010802.150424

3. Rash LD, Hodgson WC. Pharmacology and biochemistry of spider venoms. Toxicon. 2002;40: 225–254. doi:10.1016/S0041-0101(01)00199-4

4. Daltry JC, Wüster W, Thorpe RS. Diet and snake venom evolution. Nature. 1996;

5. Duda TF, Remigio EA. Variation and evolution of toxin gene expression patterns of six closely related venomous marine snails. Mol Ecol. 2008;17: 3018–32. doi:10.1111/j.1365-294X.2008.03804.x

6. Da Silva Jr. NJ, Aird SD. Prey specificity, comparative lethality and compositional differences of coral snake venoms. Comp Biochem Physiol Part C Toxicol Pharmacol. 2001;

7. Gutiérrez JM, Avila C, Camacho Z, Lomonte B. Ontogenetic changes in the venom of the snake Lachesis muta stenophrys (bushmaster) from Costa Rica. Toxicon. 1990;28: 419–426. doi:10.1016/0041-0101(90)90080-Q

8. Casewell NR, Wüster W, Vonk FJ, Harrison RA, Fry BG. Complex cocktails: the evolutionary novelty of venoms. Trends Ecol Evol. 2013;28: 219–29. doi:10.1016/j.tree.2012.10.020

9. Vonk FJ, Jackson K, Doley R, Madaras F, Mirtschin PJ, Vidal N. Snake venom: from fieldwork to the clinic. Bioessays. 2011;33: 269–279. doi:10.1002/bies.201000117

10. Olivera BM, Teichert RW. Diversity of the neurotoxic Conus peptides. Mol Interv. 2007;7: 251–260. doi:10.1124/mi.7.5.7

11. Pahari S, Bickford D, Fry BG, Kini RM. Expression pattern of three-finger toxin and phospholipase A2 genes in the venom glands of two sea snakes, Lapemis curtus and Acalyptophis peronii: comparison of evolution of these toxins in land snakes, sea kraits and sea snakes. BMC Evol Biol. 2007;7: 175. doi:10.1186/1471-2148-7-175

12. Li M, Fry BG, Kini RM. Eggs-only diet: Its implications for the toxin profile changes and ecology of the marbled sea snake (Aipysurus eydouxii). J Mol Evol. 2005;60: 81–89. doi:10.1007/s00239-004-0138-0

13. Brahma RK, McCleary RJR, Kini RM, Doley R. Venom gland transcriptomics for identifying, cataloging, and characterizing venom proteins in snakes. Toxicon. Elsevier Ltd; 2015;93: 1–10. doi:10.1016/j.toxicon.2014.10.022

14. Binford GJ. Differences in venom composition between orb-weaving and wandering Hawaiian Tetragnatha (Araneae). Biol J Linn Soc. 2001;74: 581–595. doi:10.1006/bijl.2001.0592

15. Creer S, Malhotra A, Thorpe RS, Stöcklin R, Favreau P, Chou WH. Genetic and ecological correlates of intraspecific variation in pitviper venom composition detected using matrix-assisted laser desorption time-of-flight mass spectrometry (MALDI-TOFMS) and isoelectric focusing. J Mol Evol. 2003;56: 317–329. doi:10.1007/s00239-0022403-4

16. Endean R, Rudkin C. Studies of the venoms of some Conidae. Toxicon. 1963;1: 49–64.

17. Endean R, Rudkin C. Further studies of the venoms of Conidae. Toxicon. 1965;69: 225–249. doi:10.1016/0041-0101(65)90021-8

18. Richards DP, Barlow A, Wüster W. Venom lethality and diet: Differential responses of natural prey and model organisms to the venom of the saw-scaled vipers (Echis). Toxicon. Elsevier Ltd; 2012;59: 110–116. doi:10.1016/j.toxicon.2011.10.015

19. Barlow A, Pook CE, Harrison RA, Wüster W. Coevolution of diet and prey-specific venom activity supports the role of selection in snake venom evolution. Proc Biol Sci. 2009;276: 2443–2449. doi:10.1098/rspb.2009.0048

20. Gibbs HL, Sanz L, Sovic MG, Calvete JJ. Phylogeny-based comparative analysis of venom proteome variation in a clade of rattlesnakes (Sistrurus sp.). PLoS One. 2013;8. doi:10.1371/journal.pone.0067220

21. Williams V, White J, Schwaner TD, Sparrow A. Variation in venom proteins from isolated populations of tiger snakes (Notechis ater niger, N. scutatus) in South Australia. Toxicon. 1988;26: 1067–1075. doi:10.1016/0041-0101(88)90205-X

22. Fry BG, Wüster W, Kini RM, Brusic V, Khan a., Venkataraman D, et al. Molecular evolution and phylogeny of elapid snake venom three-finger toxins. J Mol Evol. 2003;57: 110–129. doi:10.1007/s00239-003-2461-2

23. Casewell NR, Wagstaff SC, Harrison RA, Renjifo C, Wu W. Domain Loss Facilitates Accelerated Evolution and Neofunctionalization of Duplicate Snake Venom Metalloproteinase Toxin Genes. 2011;28: 2637–2649. doi:10.1093/molbev/msr091

24. Chang D, Duda TF. Extensive and continuous duplication facilitates rapid evolution and diversification of gene families. Mol Biol Evol. 2012;29: 2019–29. doi:10.1093/molbev/mss068

25. Duda TF, Palumbi SR. Molecular genetics of ecological diversification: duplication and rapid evolution of toxin genes of the venomous gastropod Conus. Proc Natl Acad Sci U S A. 1999;96: 6820–3.

26. Fry BG, Wüster W, Ryan Ramjan SF, Jackson T, Martelli P, Kini RM. Analysis of Colubroidea snake venoms by liquid chromatography with mass spectrometry: evolutionary and toxinological implications. Rapid Commun Mass Spectrom. 2003;17: 2047–2062. doi:10.1002/rcm.1148

27. Van Valen L. Morphological variation and width of ecological niche. Am Nat. 1965;99: 377–390. doi:10.2307/2678832

28. Remigio EA, Duda TF. Evolution of ecological specialization and venom of a predatory marine gastropod. Mol Ecol. 2008;17: 1156–62. doi:10.1111/j.1365-294X.2007.03627.x

29. Elliger CA, Richmond TA, Lebaric ZN, Pierce NT, Sweedler JV, Gilly WF. Diversity of conotoxin types from Conus californicus reflects a diversity of prey types and a novel evolutionary history. Toxicon. 2011;57: 311–22. doi:10.1016/j.toxicon.2010.12.008

30. Chang D, Olenzek AM, Duda Jr. TF. Effects of geographical heterogeneity in species interactions on the evolution of venom genes. Proc R Soc B. 2015; doi:10.1098/rspb.2014.1984

31. Jin A, Dutertre S, Kaas Q, Lavergne V, Kubala P, Lewis RJ, et al. Transcriptomic messiness in the venom duct of Conus miles contributes to conotoxin diversity. Mol Cell proteomics. 2013;12: 3824–33. doi:10.1074/mcp.M113.030353

32. Kaas Q, Westermann J-C, Craik DJ. Conopeptide characterization and classifications: an analysis using ConoServer. Toxicon. 2010;55: 1491–509. doi:10.1016/j.toxicon.2010.03.002

33. Li M, Fry BG, Kini RM. Putting the brakes on snake venom evolution: The unique molecular evolutionary patterns of Aipysurus eydouxii (marbled sea snake) phospholipase A2 toxins. Mol Biol Evol. 2005;22: 934–941. doi:10.1093/molbev/msi077

34. Puillandre N, Bouchet P, Duda TF, Kauferstein S, Kohn AJ, Olivera BM, et al. Molecular phylogeny and evolution of the cone snails (Gastropoda, Conoidea). Mol Phylogenet Evol. Elsevier Inc.; 2014;78: 290–303. doi:10.1016/j.ympev.2014.05.023

35. Robinson S, Norton R. Conotoxin Gene Superfamilies. Mar Drugs. 2014;12: 6058–6101. doi:10.3390/md12126058

36. Section MS, Diseases K. Conus peptides targeted to specific nicotinic acetylcholine receptor subtypes. Annu Rev Biochem. 1999;68: 59–88. doi:10.1146/annurev.pharmtox.44.101802.121622

37. Dutertre S, Jin A, Kaas Q, Jones A, Alewood PF, Lewis RJ. Deep venomics reveals the mechanism for expanded peptide diversity in cone snail venom. Mol Cell proteomics. 2013;12: 312–29. doi:10.1074/mcp.M112.021469

38. Olivera BM, Rivier J, Clark C, Ramilo CA, Corpuz GP, Abogadie FC, et al. Diversity of Conus neuropeptides. Science (80-). 1990;

39. Barghi N, Concepcion GP, Olivera BM, Lluisma AO. High conopeptide diversity in Conus tribblei revealed through analysis of venom duct transcriptome using two high-throughput sequencing platforms. Mar Biotechnol. 2014;17: 81–98. doi:10.1007/s10126-014-9595-7

40. Pi C, Liu J, Peng C, Liu Y, Jiang X, Zhao Y, et al. Diversity and evolution of conotoxins based on gene expression profiling of Conus litteratus. Genomics. 2006;88: 809–19. doi:10.1016/j.ygeno.2006.06.014

41. Barghi N, Concepcion GP, Olivera BM, Lluisma a. O. Comparison of the venom peptides and their expression in closely related Conus species: insights into adaptive post-speciation evolution of Conus exogenomes. Genome Biol Evol. 2015;7: 1797–1814. doi:10.1093/gbe/evv109

42. Duda Jr. TF, Kohn AJ, Palumbi SR. Origins of diverse feeding ecologies within Conus, a genus of venomous marine gastropods. Biol J Linn Soc. 2001;73: 391–409. doi:10.1006/bij1.2001.0544

43. Duda TF, Bolin MB, Meyer CP, Kohn AJ. Hidden diversity in a hyperdiverse gastropod genus: discovery of previously unidentified members of a Conus species complex. Mol Phylogenet Evol. 2008;49: 867–76. doi:10.1016/j.ympev.2008.08.009

44. Grabherr MG, Haas BJ, Yassour M, Levin JZ, Thompson DA, Amit I, et al. Full-length transcriptome assembly from RNA-Seq data without a reference genome. Nat Biotechnol. 2011;29: 644–52. doi:10.1038/nbt.1883

45. Altschup SF, Gish W, Pennsylvania T, Park U. Basic Local Alignment Search Tool. J Mol Biol. 1990;215: 403–410.

46. Lavergne V, Dutertre S, Jin A, Lewis RJ, Taft RJ, Alewood PF. Systematic interrogation of the Conus marmoreus venom duct transcriptome with ConoSorter reveals 158 novel conotoxins and 13 new gene superfamilies. BMC Genomics. 2013;14. doi:10.1186/1471-2164-14-708

47. Kaas Q, Yu R, Jin A-H, Dutertre S, Craik DJ. ConoServer: updated content, knowledge, and discovery tools in the conopeptide database. Nucleic Acids Res. 2012;40: D325–30. doi:10.1093/nar/gkr886

48. Lu A, Yang L, Xu S, Wang C. Various conotoxin diversifications revealed by a venomic study of Conus flavidus. Mol Cell proteomics. 2014;13: 105–18. doi:10.1074/mcp.M113.028647

49. Olivera BM, Walker C, Cartier GE, Hooper D, Santos AD, Schoenfeld R, et al. Speciation of cone snails and interspecific hyperdivergence of their venom peptides: potential evolutionary significance of introns. Ann N Y Acad Sci. 1999;870: 223–237.

50. Simakov O, Marletaz F, Cho S-J, Edsinger-Gonzales E, Havlak P, Hellsten U, et al. Insights into bilaterian evolution from three spiralian genomes. Nature. 2013;493: 526–31. doi:10.1038/nature11696

51. Kohn AJ. Tempo and mode of evolution in conidae. Malacologia. 1990;32: 55–67.

52. Sanderson MJ. r8s: inferring absolute rates of molecular evolution and divergence times in the absence of a molecular clock. Bioinformatics. 2003;19: 301–302. doi:10.1093/bioinformatics/19.2.301

53. Li B, Dewey CN. RSEM: accurate transcript quantification from RNA-Seq data with or without a reference genome. BMC Bioinformatics. 2011;12. doi:10.1186/1471-2105-12-323

54. Wagner GP, Kin K, Lynch VJ. Measurement of mRNA abundance using RNA-seq data: RPKM measure is inconsistent among samples. Theory Biosci. 2012;131: 281–5. doi:10.1007/s12064-012-0162-3

55. Schoener TW. The Anolis lizards of Bimini: resource partitioning in a complex fauna. Ecology. 1968;49: 704–726.

56. Kohn AJ. The ecology of Conus in Hawaii. Ecol Monogr. 1959;29: 47–90.

57. Kohn AJ. Food specialization in Conus in Hawaii and California. Ecology. 1966;47: 1041–1043.

58. Marsh H. Observations on the food and feeding of some vermivorous Conus on the Great Barrier Reef. The veliger. 1971;

59. Kohn AJ. Microhabitats, abundance and food of Conus on atoll reef’s in the Maldive and Chagos islands. Ecology. 1968;49: 1046–1062.

60. Kohn AJ, Nybakken JW. Ecology of Conus on eastern Indian Ocean fringing reefs: diversity of species and resource utilization. Mar Biol. 1975;29: 211–234. doi:10.1007/BF00391848

61. Kohn AJ. Abundance, diversity, and resource use in an assemblage of Conus species. Pacific Sci. 1981;34: 359–369.

62. Orme D. The caper package⁪: comparative analysis of phylogenetics and evolution in R. 2013. pp. 1–36.

63. Lavergne V, Harliwong I, Jones A, Miller D, Taft RJ, Alewood PF. Optimized deep-targeted proteotranscriptomic profiling reveals unexplored Conus toxin diversity and novel cysteine frameworks. Proc Natl Acad Sci. 2015;112. doi:10.1073/pnas.1501334112

64. Minoche AE, Dohm JC, Himmelbauer H. Evaluation of genomic high-throughput sequencing data generated on Illumina HiSeq and Genome Analyzer systems. Genome Biol. 2011;12: R112. doi:10.1186/gb-2011-12-11-r112

65. Gilles A, Meglécz E, Pech N, Ferreira S, Malausa T, Martin J-F. Accuracy and quality assessment of 454 GS-FLX Titanium pyrosequencing. BMC Genomics. BioMed Central Ltd; 2011;12: 245. doi:10.1186/1471-2164-12-245

66. Robinson SD, Safavi-Hemami H, McIntosh LD, Purcell AW, Norton RS, Papenfuss AT. Diversity of conotoxin gene superfamilies in the venomous snail, Conus victoriae. PLoS One. 2014;9: e87648. doi:10.1371/journal.pone.0087648

67. Conticello SG, Gilad Y, Avidan N, Ben-Asher E, Levy Z, Fainzilber M. Mechanisms for evolving hypervariability: the case of conopeptides. Mol Biol Evol. 2001;18: 120–31.

68. Liu Z, Li H, Liu N, Wu C, Jiang J, Yue J, et al. Diversity and evolution of conotoxins in Conus virgo, Conus eburneus, Conus imperialis and Conus marmoreus from the South China Sea. Toxicon. 2012;60: 982–989. doi:10.1016/j.toxicon.2012.06.011

69. Haney RA, Ayoub NA, Clarke TH, Hayashi CY, Garb JE. Dramatic expansion of the black widow toxin arsenal uncovered by multi-tissue transcriptomics and venom proteomics. BMC Genomics. 2014;15: 366. doi:10.1186/1471-2164-15-366

70. Casewell NR, Harrison RA, Wüster W, Wagstaff SC. Comparative venom gland transcriptome surveys of the saw-scaled vipers (Viperidae: Echis) reveal substantial intra-family gene diversity and novel venom transcripts. BMC Genomics. 2009;10: 564. doi:10.1186/1471-2164-10-564

71. Rokyta DR, Wray KP, Margres MJ. The genesis of an exceptionally lethal venom in the timber rattlesnake (Crotalus horridus) revealed through comparative venom-gland transcriptomics. BMC Genomics. 2013;14: 394. doi:10.1186/1471-2164-14-394

72. Hu H, Bandyopadhyay PK, Olivera BM, Yandell M. Characterization of the Conus bullatus genome and its venom-duct transcriptome. BMC Genomics. 2011;12. doi:10.1186/1471-2164-12-60

73. Zhang Y, Chen J, Tang X, Wang F, Jiang L, Xiong X, et al. Transcriptome analysis of the venom glands of the Chinese wolf spider Lycosa singoriensis. Zoology. 2010;113: 10–8. doi:10.1016/j.zool.2009.04.001

74. Rokyta DR, Lemmon AR, Margres MJ, Aronow K. The venom-gland transcriptome of the eastern diamondback rattlesnake (Crotalus adamanteus). BMC Genomics. 2012;13: 213.

75. Bresler M, Sheehan S, Chan AH, Song YS. Telescoper: De novo assembly of highly repetitive regions. Bioinformatics. 2012;28: 311–317. doi:10.1093/bioinformatics/bts399

76. Prjibelski AD, Vasilinetc I, Bankevich A, Gurevich A, Krivosheeva T, Nurk S, et al. ExSPAnder: A universal repeat resolver for DNA fragment assembly. Bioinformatics. 2014;30: 293–301. doi:10.1093/bioinformatics/btu266

77. Puillandre N, Koua D, Favreau P, Olivera BM, Stöcklin R. Molecular phylogeny, classification and evolution of conopeptides. J Mol Evol. 2012;74: 297–309. doi:10.1007/s00239-012-9507-2

78. Duda TF, Palumbi SR. Evolutionary diversification of multigene families: allelic selection of toxins in predatory cone snails. Mol Biol Evol. 2000;17: 1286–93.

79. Parry L, Tanner A, Vinther J. The origin of annelids. Smith A, editor. Palaeontology. 2014;57: 1091–1103. doi:10.1111/pala.12129

80. Millard EL, Daly NL, Craik DJ. Structure-activity relationships of alpha-conotoxins targeting neuronal nicotinic acetylcholine receptors. Eur J Biochem. 2004;271: 2320–6. doi:10.1111/j.1432-1033.2004.04148.x

81. Bolnick DI, Svanbäck R, Araújo MS, Persson L. Comparative support for the niche variation hypothesis that more generalized populations also are more heterogeneous. Proc Natl Acad Sci. 2007;104: 10075–10079. doi:10.1073/pnas.0703743104

82. Duda TF, Lee T. Ecological release and venom evolution of a predatory marine snail at Easter Island. PLoS One. 2009;4. doi:10.1371/journal.pone.0005558

83. Crow KD, Wagner GP. What is the role of genome duplication in the evolution of complexity and diversity? Mol Biol Evol. 2006;23: 887–92. doi:10.1093/molbev/msj083

84. Kirschner M, Gerhart J. Evolvability. Proc Natl Acad Sci U S A. 1998;95: 8420–8427.

85. Pyron RA, Burbrink FT. EXTINCTION, ECOLOGICAL OPPORTUNITY, AND THE ORIGINS OF GLOBAL SNAKE DIVERSITY. 2011; 163–178. doi:10.5061/dryad.63kf4

86. Dutertre S, Jin A-H, Vetter I, Hamilton B, Sunagar K, Lavergne V, et al. Evolution of separate predation-and defence-evoked venoms in carnivorous cone snails. Nat Commun. 2014;5: 3521. doi:10.1038/ncomms4521

87. Jin A, Israel MR, Inserra MC, Smith JJ, Lewis RJ, Alewood PF, et al. delta-Conotoxin SuVIA suggests an evolutionary link between ancestral predator defence and the origin of fish-hunting behaviour in carnivorous cone snails. Proc R Soc B. 2015;282. doi:10.1098/rspb.2015.0817

88. Aman JW, Imperial JS, Ueberheide B, Zhang M-M, Aguilar M, Taylor D, et al. Insights into the origins of fish hunting in venomous cone snails from studies of *Conus tessulatus*. Proc Natl Acad Sci. 2015;112: 5087–5092. doi:10.1073/pnas.1424435112

89. Hu H, Bandyopadhyay PK, Olivera BM, Yandell M. Elucidation of the molecular envenomation strategy of the cone snail Conus geographus through transcriptome sequencing of its venom duct. BMC Genomics. 2012;13. doi:10.1186/1471-2164-13-284

90. Bolger AM, Lohse M, Usadel B. Trimmomatic: a flexible trimmer for Illumina sequence data. Bioinformatics (Oxford, England). 2014. pp. 1–7. doi:10.1093/bioinformatics/btu170

91. Magoč T, Salzberg SL. FLASH: fast length adjustment of short reads to improve genome assemblies. Bioinformatics. 2011;27: 2957–63. doi:10.1093/bioinformatics/btr507

92. Langmead B, Salzberg SL. Fast gapped-read alignment with Bowtie 2. Nat Methods. 2012;9: 357–9. doi:10.1038/nmeth.1923

93. Safavi-Hemami H, Gajewiak J, Karanth S, Robinson SD, Ueberheide B, Douglass AD, et al. Specialized insulin is used for chemical warfare by fish-hunting cone snails. Proc Natl Acad Sci. 2015;112: 1743–1748. doi:10.1073/pnas.1423857112

94. Huang X, Madan A. CAP3: A DNA Sequence Assembly Program. Genome Res. 1999;9: 868–877.

95. Katoh K, Kuma K-I, Toh H, Miyata T. MAFFT version 5: Improvement in accuracy of multiple sequence alignment. Nucleic Acids Res. 2005;33: 511–518. doi:10.1093/nar/gki198

